# Physical models of spatial genome organization

**DOI:** 10.1101/089755

**Authors:** S.V. Slanina, A.V. Aleshchenko, Y.A. Eidelman, S.G. Andreev

## Abstract

Remodeling of nuclear organization occurs during normal cell development, differentiation and cancer. One of the biggest gaps of knowledge remains how to link the information on chromatin and chromosome structural organization with genes activity. In this paper we introduce some physical ideas and a general computational method demonstrating how genome 3D architecture and its remodeling can be quantitatively modeled. We study a hypothetical scenario of alterations of chromosome territories positioning in the course of cell proliferation. On this basis we obtain quantitative information about chromosomal contacts in the nucleus. We predict changes of radial distributions of contacts between chromosomal megabase domains during proliferation. The proposed modeling approach may be helpful in integrating experimental data on nuclear reorganization associated with normal development and with various diseases. This predictive modeling may find applications in genome research of normal and cancer cells, stem cell biology, biology of aging, *etc*.

## INTRODUCTION

Spatial organization of genome plays an important role in cell functioning. The chromosome and gene positions in a nucleus plays a functional role in regulating their expression in mammalian cells [Chandramouly et al. 2007, Harewood et al. 2010]. Changes of gene expression may be a consequence of alterations in normal chromosome territory positioning. These changes are often seen during tumorigenesis-associated genome instability. Large scale and locus-specific reorganization of the genome during differentiation and dramatic alterations of nuclear organization associated with cancer development are well documented [Chandramouly et al. 2007, Meaburn 2016]. Experimental and modeling data suggest very complex relationships between structural and functional organization of the genome [Dekker 2014, Nowotny et al. 2016].

The aim of this work is to develop physics-based modeling approach in order to study spatial organization of genome, first of all, large scale structure of chromosomes in the nuclei. We show that proposed changes in chromosome positioning during proliferation result in significant alteration of distribution of chromosomal subunits or domains and their contacts in the nucleus. We demonstrate how these changes may be quantified by predicting different probabilistic distributions. This approach can be applied in the future to the analysis of structural organization of cells of different types, normal vs malignant cells, senescent cells, *etc*.

## METHODS

Multiscale modeling of large-scale spatial organization of all 46 chromosomes in an interphase (G1) human cell nucleus is performed by extension of earlier methods [Eidelman et al. 2006, 2012]. Each chromosome is represented as a coarse-grained chain of semi-rigid elements corresponding to megabase sized chromosomal subunits, or domains [Eidelman and Andreev 2002, Eidelman et al. 2012]. G1 chromosome structures are established in the course of decondensation of mitotic chromosomes. Subunits within a chromosome interact through a Lennard-Jones potential, subunits from different chromosomes through a truncated Lennard-Jones potential with only the excluded volume component. A Metropolis method with the dynamic variant of Monte Carlo (MC) technique is used, at each MC step positions of elements are randomly changed [Eidelman et al. 2006, 2012].

Proposed alteration of chromosome repositioning in the course of proliferation is modeled according the general algorithm described below. In each time interval (which can correspond to one or several cell divisions) several or all chromosomes change radial distributions of their centers of mass in such a way that initially peripheral chromosomes gradually change positions toward the interior of nucleus and *vice versa*. We generate 7 nucleus models in total corresponding to 7 (arbitrary) time intervals. The changes of chromosomal contacts pattern due to chromosome repositioning are calculated.

## RESULTS AND DISCUSSION

In the present work nuclear organization is characterized by radial distribution of all domains of all chromosomes. During proliferation the distribution is changed resulting in reorganization of the pattern of chromosomal contacts.

On the first step seven nucleus models are built which reflect seven time intervals of proliferation. Within each interval radial distribution of chromosomes is assumed to be unchanged. The first model corresponding to initial proliferation time, “t=0” is built to agree with the experimental radial distribution of chromosome territories for human lymphocytes [Boyle et al. 2001]. For each time interval several characteristics are determined: radial distributions of domain density and of pairwise contacts between chromosomal domains, mean number of interchromosomal contacts. The calculated changes in domain density radial distribution in the course of proliferation are presented in Fig.1.

**Fig.1.**
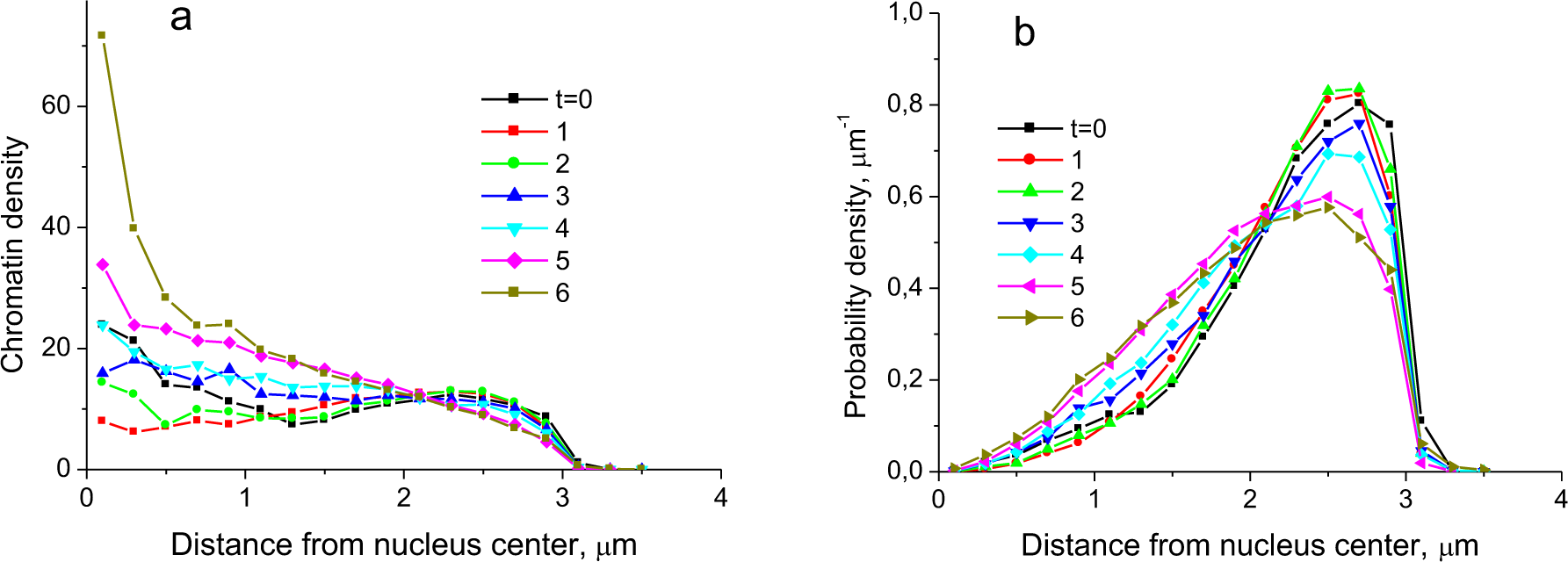
Change of radial distribution of chromatin in nuclei during proliferation. Time t (discrete number, arbitrary units) is time of proliferation. Within each time nuclear distribution of chromosome territories remains unchanged. In the model nuclear positioning is changed between different “times” abruptly by setting discrete change of spatial distributions of chromosome territories in the nuclei. (a) - radial dependence of density of chromosomal subunits, or megabase domains in a nucleus. (b) - probability density of location of chromosome subunits in a spherical layer at distance r, r+δr from nucleus center.

Fig.1b shows that at the beginning of proliferation interphase chromatin is distributed preferentially at periphery of the nucleus (black curve). This corresponds to relatively uniform chromatin density, Fig.1a. During cell proliferation chromatin is shifted to more central regions of the nucleus, its density becomes more nonhomogeneous. It increases at distances closer than 2 μm to the nuclear center, i.e. up to 2/3 of nucleus radius, and decreases at large distances.

As an output results we get a set of 3D models of seven different populations of nuclei. They contain the following information: 3D conformations and radial distributions of positions of all chromosomes in nuclei, distributions of chromatin density. On this basis we calculate number and spatial distributions of interchromosomal contacts between any chromosomes as a function of proliferation time.

During proliferation chromosome territories are shifted generally towards the nucleus interior and contacts between chromosomal domains also become more central, Fig.2a. Moreover, the number of interchromosomal contacts increases by 30% at the end of proliferation with respect to the nonproliferating initial state, Fig.2b.

**Fig.2.**
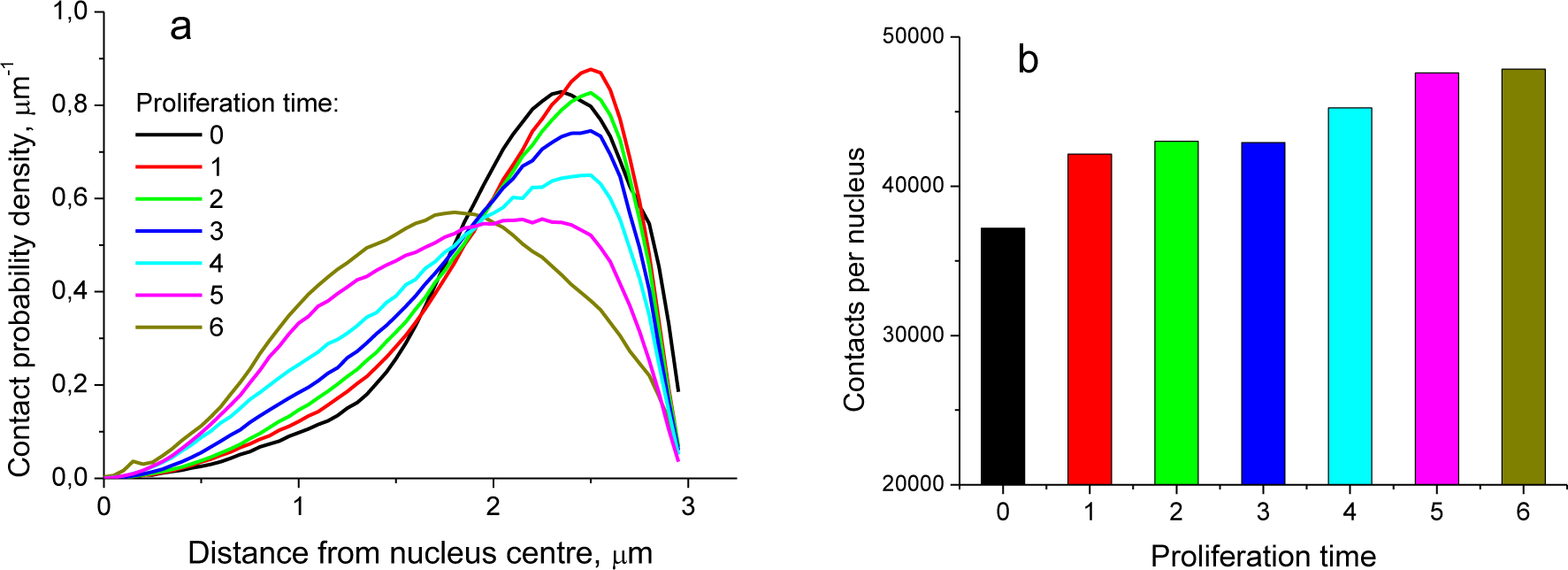
Changes of distribution of interchromosomal contacts in the course of proliferation. (a) - radial distribution. (b) - number of contacts per nucleus. Different colors on the right mean the same different time intervals as on the left.

We generate the 3D structure for one initial nucleus (at t=0) and follow it during proliferation, Fig.3. For viewing convenience relative positioning of only two chromosomes, 13 and 22, is tracked. Other chromosomes and their domains in the nucleus are drawn for simplicity in semitransparent blue balls.

**Fig.3.**
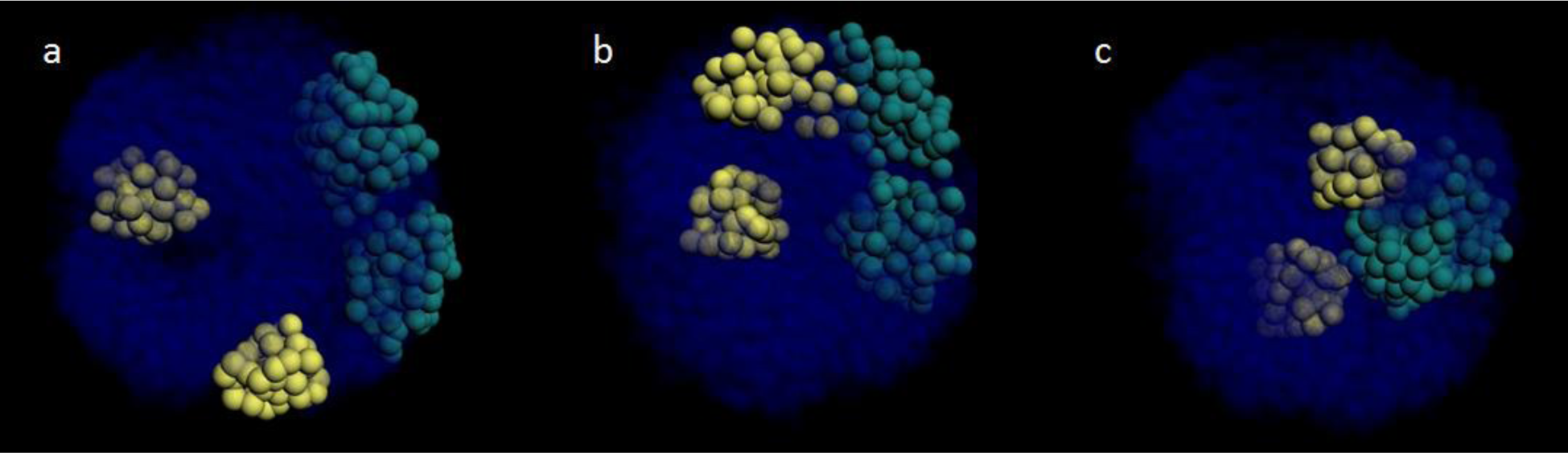
Visualization of chromosomal contacts for different times of proliferation of the single nucleus. (a) – initial time. (b) – intermediate time. (c) – late time. Colored balls represent 1 Mbp chromosomal domains. Two chromosomes 13 and 21 are in different colors, chromosome 13 in dark cyan, chromosome 21 in yellow. Semitransparent blue balls are domains from all other chromosomes.

One can observe in 3D how position of different chromosome territories as well as chromosomal domains is altered during proliferation and how relative proximity of chromosomal and genetic loci is changed. Some loci, or megabase domains, become juxtaposed in time. We speculate that this type of mechanisms may underlie the formation of cancer specific translocations *in vivo*.

Chromatin repositioning including relocation of single or multiple loci in different parts of chromosomes has been proposed as a cell response to radiation exposure [Andreev and Spitkovsky 1987]. This proposal was confirmed recently. The evidence of DNA damage dependent large-scale spatial reorganization of chromosomes has been documented which might serve as an important aspect of cellular damage response [Mehta et al. 2013].

Interphase chromosome positioning including chromosome intermingling, number of chromosomal contacts and centers, active/inactive chromatin structure and dynamics, *etc*. might contribute to chromosomal aberration formation [Eidelman and Andreev 2011]. Impact of interphase chromosome positioning on the spectrum of radiation-induced chromosomal aberrations was shown [Boei et al. 2006].

The modeling technique presented demonstrates ability to perform comprehensive mapping of spatial positioning patterns of chromosomes, contacts and specific genomic loci in nuclei at different times of proliferation. Repositioning of chromosomes in the nucleus may be associated with changes in gene expression. According to the model, some chromosomes are relocated during proliferation toward periphery and some toward nucleus interior. The placement of genes at new environment position due to their relocation relative to the center/periphery of the nucleus can alter their expression, for example, due to change in relative proximity with transcription factories. It has been observed that gene expression can depend on their movement toward the nuclear periphery or interior [Harewood et al. 2010]. The modeled hypothetical scenario may resemble the nuclear reprogramming process or adaptation of nuclear architecture to the prolonged cell culturing, or alteration of nuclear organization during immortalization and neoplastic transformation. Genome structure-oriented quantitative modeling may help to understand how cancer translocations are formed during aging or under environmental DNA-damaging factors.

In summary, the polymer physics-based multilevel modeling technique for studying spatial organization of genome is presented. It allows to predict spatial distributions of chromosomal contacts in the nuclei of proliferating cells. It takes into account alterations of spatial positioning of chromosomes resulting in changes of interchromosomal contacts pattern. The results obtained provide a basis for future studies of genome structure and reorganization in various areas: radiobiology (e.g. predicting delayed effects of irradiation like genomic instability), cancer research, stem cell biology, *etc*.

## ACKNOWLEDGEMENTS

The present work is supported by Russian Foundation for Basic Research grant 14-01-00825 to S.A.

